# Multiscale Entropy of Resting-State fMRI Signals Reveals Differences in Brain Complexity in Autism

**DOI:** 10.1101/2025.05.22.655518

**Authors:** Wenyi Xiao, Myles Jones

## Abstract

**Background:** Atypical intrinsic brain activity has been widely observed in autism spectrum disorder (ASD), yet the temporal complexity of these neural signals remains underexplored. This study aimed to characterise differences in resting-state brain signal complexity between individuals with ASD and neurotypical controls using multiscale entropy (MSE).

**Methods:** Resting-state fMRI data were obtained from the Autism Brain Imaging Data Exchange I (ABIDE I), a large multi-site dataset including 397 participants (179 with ASC, 218 neurotypical controls; ASD: Mean age 16.43 years old, SD 7.17 years old; CON: Mean age 15.75 years old, SD 5.67 years old). Voxel-wise multiscale entropy (MSE) features were extracted across multiple temporal scales. Group comparisons were conducted using voxel-wise t-tests and mixed-effects models to identify region- and scale-specific alterations in brain signal complexity.

**Results:** Individuals with ASD showed reduced MSE in prefrontal regions at coarser time scales and elevated MSE in posterior midline regions, including the posterior cingulate cortex and precuneus, at finer scales, followed by a decline across coarser scales. This pattern suggests a shift toward uncorrelated randomness in posterior regions and reduced long-range complexity in frontal areas. No significant associations were found between MSE features and ADOS scores.

**Conclusions:** These findings reveal spatially and temporally specific alterations in brain signal complexity in ASD, particularly within the default mode network. Multiscale entropy provides a complementary approach to traditional connectivity and single-scale entropy analyses, offering novel insights into the organisation of intrinsic brain activity in neurodevelopmental conditions.

## Introduction

Resting-state functional magnetic resonance imaging (rs-fMRI) is a widely employed neuroimaging technique for examining spontaneous fluctuations in brain activity in the absence of externally imposed tasks. By measuring blood oxygen level-dependent (BOLD) signal variations, rs-fMRI enables the characterisation of intrinsic functional organisation and temporal dynamics of the brain (Lv et al., 2018). In the context of autism spectrum disorder (ASD), a neurodevelopmental condition characterised by atypical socio-communicative behaviours and restricted, repetitive patterns of activity (American Psychiatric Association, 2013), rs-fMRI studies have revealed disruptions in local and distributed brain dynamics (Hull et al., 2017). However, many of these studies rely on linear analyses that may not fully capture the complexity and nonlinear temporal structure of spontaneous brain activity.

To address this limitation, nonlinear signal processing approaches have been increasingly adopted to examine intrinsic neural dynamics, particularly those that quantify the temporal complexity of brain signals. Sample entropy and multiscale sample entropy (MSE) are among the most prominent methods in this domain. Sample entropy quantifies the regularity of a time series by evaluating the likelihood that sequences of a certain length will be repeated within the signal, with lower entropy indicating greater predictability and reduced signal complexity (Richman & Moorman, 2000; Delgado-Bonal & Marshak, 2019). MSE extends this concept by assessing entropy across multiple temporal scales using a coarse-graining procedure (Costa et al., 2002; Costa et al., 2005), thereby capturing hierarchical organisation within the signal. This is particularly important in biological systems, where relevant information may be distributed across time scales due to the nested nature of physiological processes.

In ASD, studies applying entropy-based methods to rs-fMRI have identified alterations in the complexity of brain signals across several regions, including the prefrontal cortex and sensory networks (Zhang et al., 2020; Maximo et al., 2021). Reductions in entropy have been interpreted as reflecting a loss of dynamic range or decreased flexibility in spontaneous neural activity, consistent with theoretical models suggesting that less complex brain signals are associated with reduced adaptability and impaired system functioning (Goldberger et al., 2002; Ho et al., 1997). Importantly, such reductions in entropy have also been linked to clinical symptomatology, including social communication difficulties and restricted interests, as captured by measures such as the Autism Diagnostic Observation Schedule (ADOS) and the Social Responsiveness Scale (SRS) (Easson & McIntosh, 2019; Zhang et al., 2020).

Although electroencephalography (EEG) research has provided robust evidence for altered neural complexity in ASD, with multiple studies reporting atypical sample entropy and multiscale entropy patterns in resting-state data (Bosl et al., 2011; Catarino et al., 2011; Hadoush et al., 2019), it is important to highlight the unique strengths of fMRI for studying spatially distributed brain function. While fMRI offers lower temporal resolution compared to EEG, it provides much finer spatial resolution, enabling the examination of complexity within and between large-scale neural networks. Entropy-based metrics derived from rs-fMRI thus complement EEG findings by revealing not only *how* brain activity varies over time, but also *where* in the brain these variations occur. Despite the relatively limited number of fMRI studies in this area, existing work has demonstrated that multiscale entropy can be reliably estimated from BOLD signals using conservative parameters (e.g., scale factors up to 5, m = 2, r = 0.6) (Wang et al., 2014; Nezafati et al., 2020), and that these measures are sensitive to diagnostic group differences in ASD. These findings highlight the potential of MSE as a spatially resolved marker of intrinsic brain complexity, capable of characterising disrupted temporal organisation in ASD even with the lower sampling frequency of fmri.

Nonetheless, the application of MSE to rs-fMRI is not without challenges. A key limitation stems from the relatively short duration of fMRI recordings, typically consisting of only 100–200 time points (with TR ≈ 2 s), which constrains the maximum number of scales over which entropy can be reliably computed. Coarse-graining can introduce additional noise into the signal, particularly at higher scales, potentially reducing estimation stability (Courtiol et al., 2016; Sokunbi, 2014). However, studies have shown that coarse-graining also removes short-range linear dependencies and can reveal more meaningful nonlinear structure by emphasising slower fluctuations (Govindan et al., 2007; Vakorin et al., 2011). These findings suggest that MSE remains a valid and informative approach when applied with appropriate methodological considerations.

To systematically investigate neural signal complexity in ASD, the present study adopts a hypothesis-free, data-driven analytical framework. This framework is modelled on neurophysiological biomarker pipelines (Heunis et al., 2016), comprising four stages: data acquisition, uniform preprocessing, feature extraction via voxel-wise multiscale sample entropy estimation, and statistical inference. By estimating entropy across multiple temporal scales, this approach captures hierarchical temporal structure within the BOLD signal and allows for the spatial localisation of group-level differences in signal complexity.

The decision to employ a voxel-wise, data-driven method was informed by inconsistencies in the existing literature. Prior studies have reported both increased and decreased entropy in ASD, with effects varying across brain regions and methodological approaches (Maximo et al., 2021; Zhang et al., 2020), while others have found no significant group differences (Easson & McIntosh, 2019). These discrepancies may arise from limited statistical power, differences in scale parameters, or reliance on predefined regions of interest. To address these limitations, we analyse a large, standardised subsample from the Autism Brain Imaging Data Exchange I (ABIDE I; Di Martino et al., 2014), restricted to eyes-open acquisitions with a consistent sampling rate (TR = 2000 ms), thereby enhancing the comparability and robustness of entropy estimates.

The objective of this study is to assess whether multiscale entropy can reliably detect alterations in spontaneous brain dynamics in individuals with ASD. Through whole-brain voxel-wise comparisons, we aim to characterise the spatial distribution of signal complexity differences between ASD and neurotypical control groups.

## Method

### Participants and Imaging Data

Resting-state functional magnetic resonance imaging (rs-fMRI) data were obtained from the Autism Brain Imaging Data Exchange I (ABIDE I; Di Martino et al., 2014), a publicly available repository aggregating multi-site neuroimaging data from individuals with autism spectrum disorder (ASD) and neurotypical controls. All data used in this study were obtained from the ABIDE dataset hosted by the Preprocessed Connectomes Project (http://preprocessed-connectomes-project.org/abide/), where they had already been preprocessed and made publicly available by the Neuro Bureau Preprocessing Initiative (Craddock et al., 2013). Preprocessing was conducted using the Data Processing Assistant for Resting-State fMRI (DPARSF; Chao-Gan & Yu-Feng, 2010), based on Statistical Parametric Mapping (SPM; Friston et al., 2007) and the DPABI toolbox (Yan et al., 2016). All original data acquisition was approved by the local ethics committees of contributing sites, and analyses were conducted on anonymised datasets.

From the full ABIDE I sample of 1,112 participants (539 with autism spectrum disorder [ASD] and 573 neurotypical controls), we selected participants based on both data quality and acquisition compatibility with multiscale entropy analysis. Preprocessed data were first filtered to include only those with mean framewise displacement (FD) < 0.2 mm, resulting in 884 individuals (408 ASD, 476 CON). Given the sensitivity of entropy estimation to sampling frequency, only participants with a consistent repetition time (TR = 2000 ms; 0.5 Hz sampling rate) were included. This decision was based on considerations regarding the stability and interpretability of multiscale entropy (MSE) measures across sites with different temporal resolutions. Data were drawn from seven ABIDE I sites that met this criterion: New York University Langone Medical Center (NYU), San Diego State University (SDSU), Stanford University, Trinity Centre for Health Sciences (Trinity), University of Michigan (UM), University of Southern Mississippi (USM), and Yale University. Because signal amplitude characteristics can vary across scanners and acquisition protocols, voxel-wise time series were normalised to unit variance prior to entropy estimation to minimise the influence of site-level variability on entropy values. A full rationale for site selection and signal normalisation is provided in the Supplementary Methods. This yielded 506 participants (231 ASD, 275 CON) from the included sites.

Further motion control was applied using site-specific thresholds (Supplementary Table 2). Subjects with mean FD exceeding the site-specific group mean plus two standard deviations were excluded, yielding a final sample of 500 participants (226 ASD, 274 CON).

To reduce state-related variability in BOLD signal, only eyes-open resting-state scans were analysed, yielding a final sample of 397 participants (179 with ASD, 218 neurotypical controls). A detailed flowchart of the selection procedure is provided in Supplementary Figure 1. The final sample ranged in age from 6.47 to 50.22 years (ASD: Mean 16.43 years old, SD 7.17 years old; CON: Mean 15.75 years old, SD 5.67 years old), with no significant age difference between groups (*t* = 1.04, *p* = .299). Age distributions are shown in Supplementary Figure 2, and group characteristics are summarised in Table 1.

**Table 1.** Phenotypic characteristics of the final sample (eyes-open condition, 0.5 Hz sampling rate) after quality control.

| Group | Age | Gender<br>(M/F) | FIQ | VIQ | PIQ |
| --- | --- | --- | --- | --- | --- |
| ASD | 16.43(7.17) | 157/22 | 104.16(18.22) | 103.30(20.17) | 104.40 (18.92) |
| CON | 15.75(5.67) | 165/53 | 110.87(13.49) | 112.11 (13.77) | 107.53 (13.84) |

**Note.** ASD = Autism Spectrum Disorder; CON = Neurotypical Controls.
FIQ = Full-Scale IQ; VIQ = Verbal IQ; PIQ = Performance IQ.
Values represent mean (standard deviation).

### fMRI Acquisition and Preprocessing

Resting-state fMRI data were acquired at seven ABIDE I sites using echo-planar imaging sequences with a consistent repetition time (TR) of 2000 ms, corresponding to a sampling rate of 0.5 Hz. While TR was matched across all selected datasets, acquisition parameters such as voxel dimensions, slice thickness, echo time (TE), and scan duration varied slightly by site. Full acquisition parameters are provided in Supplementary Table 1. The number of volumes ranged from 150 to 300, with voxel sizes typically between 3.0 and 3.4 mm in-plane, and slice thicknesses ranging from 3.0 mm to 4.5 mm. These site-specific protocols reflect harmonised but non-identical implementations across imaging centres.

Preprocessing was carried out by the Neuro Bureau and obtained from the ABIDE Preprocessed repository (http://preprocessed-connectomes-project.org/abide/dparsf.html; Craddock et al., 2013). The pipeline used the Data Processing Assistant for Resting-State fMRI (DPARSF; Chao-Gan & Yu-Feng, 2010), implemented on top of Statistical Parametric Mapping (SPM; Friston et al., 2007) and the DPABI toolbox (Yan et al., 2016). Key preprocessing steps included discarding the first 10 volumes, slice timing correction, realignment using a six-parameter rigid body transformation, coregistration of T1-weighted anatomical images, segmentation into grey matter, white matter, and cerebrospinal fluid (Ashburner & Friston, 2005), and spatial normalisation to the MNI152 template using DARTEL (Ashburner, 2007). Normalised functional images were resampled to 3 × 3 × 3 mm resolution and smoothed using a 6 mm full-width at half-maximum Gaussian kernel.

To mitigate motion-related confounds, 24 motion parameters were regressed out using the Friston model (Friston et al., 1998). Signals from white matter and cerebrospinal fluid, as well as linear and quadratic trends, were also removed. We used the “filt_noglobal” preprocessing strategy, which applies band-pass filtering (0.01–0.1 Hz) but does not include global signal regression.

All images were reoriented to the anterior commissure using standard reorientation matrices (https://www.rfmri.org/DownloadedReorientMats) prior to preprocessing.

### Head Motion Quality Control

Framewise displacement (FD) was calculated using Jenkinson’s method (Jenkinson et al., 2002) to quantify participant motion across the scan. During the initial inclusion phase, only participants with a mean FD below 0.2 mm were retained. To further reduce motion-related artefacts, additional site-specific thresholds were applied. At each site, the group mean FD was computed and participants with values exceeding the mean plus two standard deviations were excluded. These thresholds were calculated separately for the ASD and control groups to account for site-specific variation and to avoid disproportionately excluding ASD participants due to higher motion, thereby helping to maintain group balance and representativeness (Satterthwaite et al., 2012).

Final sample sizes and motion-related statistics per site are reported in Supplementary Table 2. Following exclusion of high-motion participants, group differences in FD were tested using independent samples t-tests within each site. Most sites showed no significant group differences (SDSU: *t* = –0.084, *p* = .934; Stanford: *t* = –0.395, *p* = .696; UM_2: *t* = 0.300, *p* = .766; USM: *t* = –0.006, *p* = .995; Yale: *t* = 1.189, *p* = .241). However, significant differences were observed at a few sites, including NYU (*t* = 3.471, *p* < .01), Trinity (*t* = 2.918, *p* < .01), and UM_1 (*t* = 3.123, *p* < .01). Across the final sample, all participants fell well within established thresholds for resting-state fMRI, supporting the validity of the comparisons and minimising residual motion confounds.

### Statistical analysis

To examine group-level differences in brain signal complexity between individuals with ASD and neurotypical controls, we performed voxel-wise statistical analyses on multiscale sample entropy (MSE) brain maps derived from resting-state fMRI data. The analysis pipeline consisted of data preprocessing, voxel-wise entropy estimation, and statistical inference using a non-parametric, cluster-based framework (Figure 1), all conducted in standard MNI space following spatial normalisation of each data.

**Figure 1.**
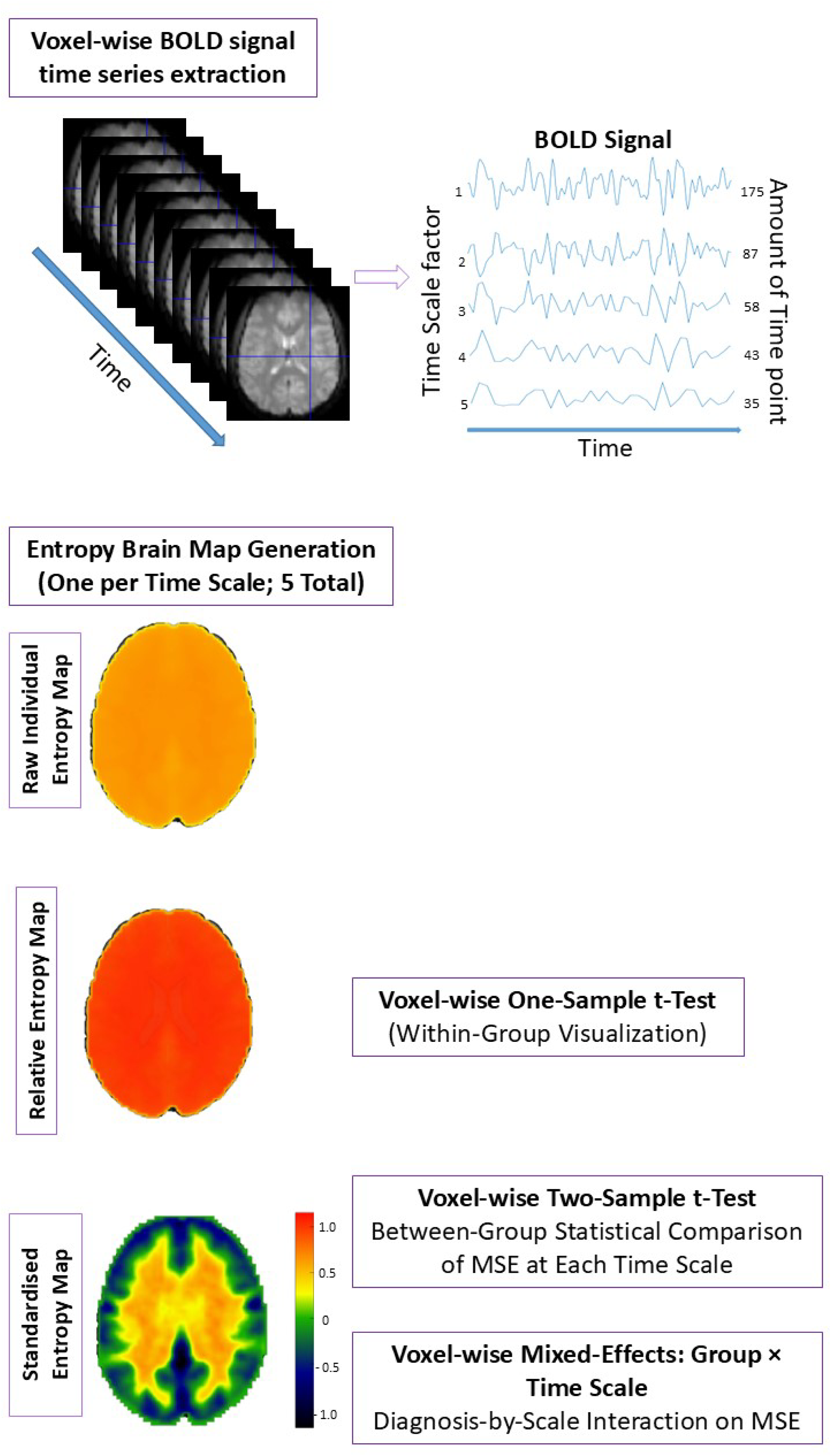
Voxel-wise BOLD time series were extracted and coarse-grained across five temporal scales (top). For each scale, a whole-brain entropy map was generated, resulting in five entropy maps per participant (middle left). Raw individual entropy maps were transformed into relative and standardised maps to support within- and between-group comparisons. Voxel-wise statistical analyses included: (i) one-sample t-tests to visualise within-group entropy distributions, (ii) two-sample t-tests to assess between-group differences at each time scale, and (iii) mixed-effects modelling to evaluate Group × Time Scale interactions (bottom right).

Voxel-wise MSE brain maps were computed for each participant across five temporal scales (τ = 1–5), using sample entropy parameters m = 2 and r = 0.6 × standard deviation (Richman and Moorman 2000). At each scale, coarse-grained time series were generated by averaging non-overlapping segments of the original signal, and sample entropy was then computed using a MATLAB implementation adapted from Azami et al. (2017). Prior to entropy estimation, voxel-wise BOLD time series were normalised to unit standard deviation to reduce variability in signal amplitude across sites.

Voxel-wise BOLD time series were extracted in MNI space for each participant, and MSE was then computed at each voxel across five time scales. The resulting entropy maps were smoothed with a 6 mm full-width at half-maximum (FWHM) Gaussian kernel to prepare them for statistical comparison. All steps were implemented in MATLAB using custom scripts and functions from the SPM and DPABI toolboxes. Full implementation details and statistical analyses are provided in the Supplementary Materials. To ensure the validity of MSE applied to fMRI data, we conducted a control analysis using phase-shuffled surrogate time series to confirm that MSE is sensitive to nonlinear signal characteristics. These validation results are reported in the Supplementary Materials (see Supplementary Figure 3).Code for entropy estimation and map generation is available at https://github.com/wenyixiao0058/fmristudy.

To characterise local entropy in relation to overall brain-wide complexity, voxel-wise MSE values were normalised by each participant’s global mean entropy, resulting in relative entropy maps referred to as *smentropy* (Li et al., 2021). This normalisation allows regional entropy values to be expressed as a proportion of the individual’s global mean, facilitating group-level comparisons of spatial entropy patterns. Global means were calculated within a group-level brain mask that included voxels present in at least 90% of participants. Global means were computed within a group-level brain mask that included voxels present in at least 90% of participants, ensuring consistent spatial sampling. The resulting *smentropy* maps allowed for regionally specific comparisons across participants while accounting for individual differences in overall signal complexity.

All voxel-wise statistical maps were corrected for multiple comparisons using threshold-free cluster enhancement (TFCE), implemented in the PALM (Permutation Analysis of Linear Models) toolbox with 5,000 permutations and no acceleration, as recommended for spatial inference (Winkler et al., 2016). TFCE avoids arbitrary cluster-forming thresholds and maintains sensitivity to spatially extended effects while controlling the family-wise error rate (Smith & Nichols, 2009).

To assess within-group spatial patterns of relative entropy, voxel-wise one-sample t-tests were performed separately for the ASD and CON groups, testing whether *smentropy* at each voxel significantly deviated from 1. A value of 1 indicates that a voxel’s entropy matches the individual’s global mean, while values above or below 1 indicate higher or lower local entropy, respectively. These tests were performed using DPABI functions and corrected using TFCE. *Neuroimage*, *44*(1), 83–98. https://doi.org/10.1016/j.neuroimage.2008.03.061. The resulting t-statistics were converted to Z-maps and visualised with MRIcroGL (Rorden & Brett, 2000). Additionally, no significant differences in global mean entropy were found between groups at any scale (Scale 1: *t* = –1.111, *p* = .267; Scale 2: *t* = –2.532, *p* = .012; Scale 3: *t* = –1.691, *p* = .092; Scale 4: *t* = –0.925, *p* = .355; Scale 5: *t* = –0.748, *p* = .455), supporting the use of normalised entropy for group comparisons.

Group-level voxel-wise analyses were conducted to examine differences in brain signal complexity between ASD and control groups across five temporal scales. Prior to statistical testing, sample entropy maps were standardised within each participant to account for inter-individual variability in global signal complexity. Specifically, voxel-wise entropy values were transformed to z-scores by subtracting the global mean and dividing by the standard deviation within each participant’s brain mask, excluding non-brain voxels. The resulting standardised entropy maps were used in all group-level statistical analyses, including two-sample *t*-tests at each scale and mixed-effects models assessing diagnosis-by-scale interactions. This normalisation approach aligns with established practices in resting-state fMRI research using the DPABI toolbox (Xiong et al., 2024). Full details of the group-level statistical procedures are provided in the Supplementary Methods.

### Exploratory Correlation with Clinical Scores

To assess the potential clinical relevance of neurophysiological markers, exploratory analyses were conducted to examine the association between sample entropy values and autism symptom severity. Pearson’s correlations were computed between ADOS total scores and mean standardised entropy extracted from the significant clusters identified in the voxel-wise two-sample *t*-tests. This analysis was performed on a subset of 108 ASC participants from the NYU, SDSU, and USM sites, for whom ADOS scores were available. Further details are provided in the Supplementary Methods and Supplementary Table 3 and 4.

## Results

### Voxel-wise one sample t test result

Spatial patterns of relative entropy (*smentropy*) within the ASD and control groups are shown in Figure 2. Voxel-wise one-sample *t*-tests were performed separately for each group at each temporal scale, without covariates, to test whether *smentropy* at each voxel significantly differed from 1. Results were corrected using threshold-free cluster enhancement (TFCE) and transformed into *Z*-maps for visualisation.

**Figure 2.**
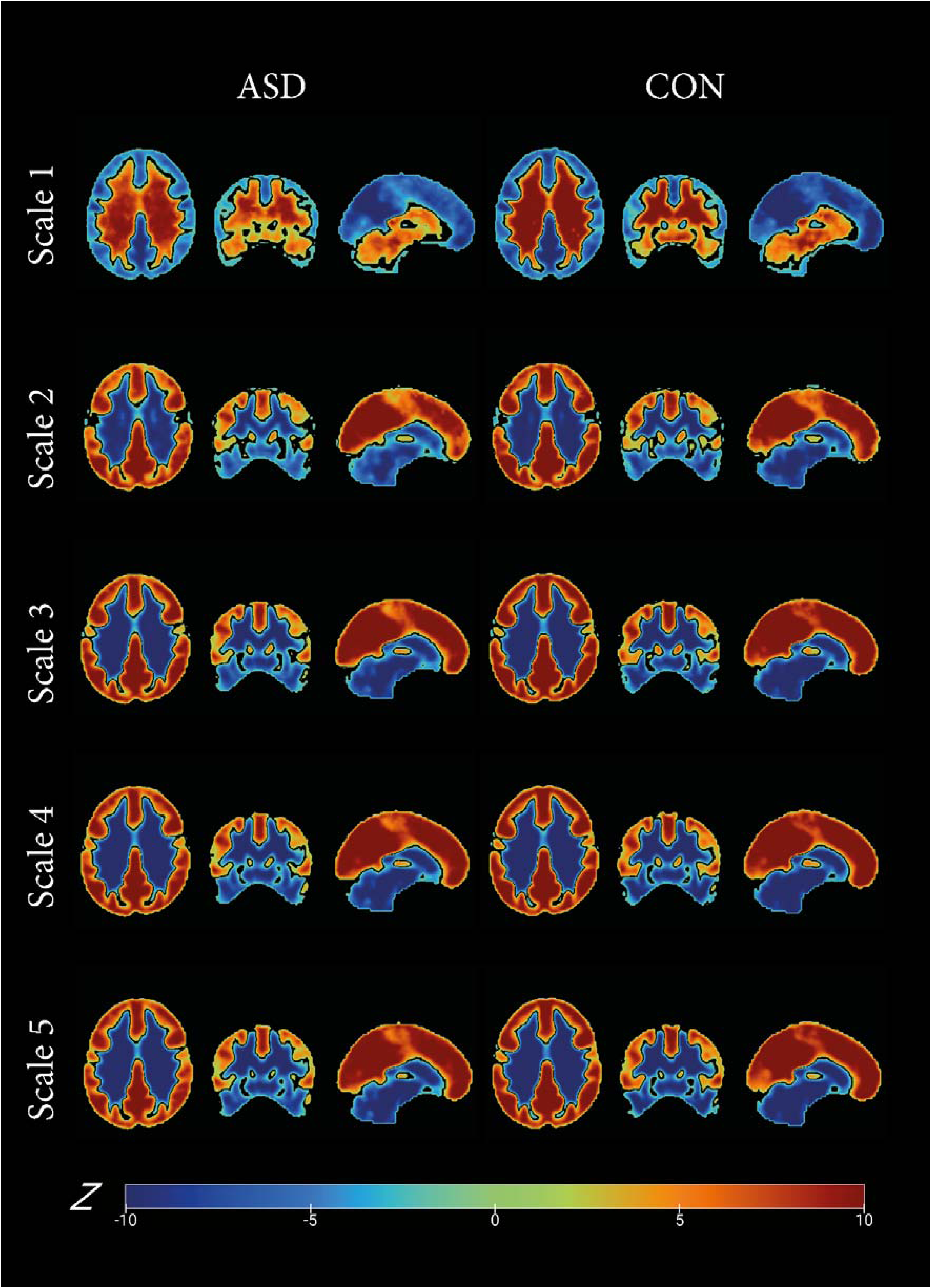
Within-group brain maps of voxel-wise multiscale sample entropy across five temporal scales in individuals with autism spectrum condition (ASD; left) and neurotypical controls (right). Each row shows entropy maps at a different time scale (1 to 5), computed from resting-state fMRI BOLD signals. Warmer colours (e.g., red/yellow) indicate higher-than-average entropy values relative to 1 (i.e., the baseline for relative entropy), while cooler colours indicate lower values. Maps reflect results from one-sample t-tests within each group and scale, corrected using threshold-free cluster enhancement (TFCE) with voxel-level permutation testing at *p* < .05. Z-statistics are displayed on a symmetric colour scale. Note: These maps are presented for visualisation purposes only and do not reflect between-group statistical comparisons.

Visual inspection of the resulting maps reveals distinct tissue-dependent patterns. At Scale 1, higher relative entropy is primarily observed in grey matter regions, while white matter exhibits lower entropy. From Scale 2 onward, this pattern reverses: white matter regions show increased relative entropy, while grey matter becomes comparatively lower. This reversal becomes more pronounced at higher scales, indicating a shift in the spatial distribution of signal complexity as the temporal scale increases.

### Voxel-wise two sample t test result: Group Level Statistical Analysis

Figure 3 illustrates the results of voxel-wise two-sample *t*-tests comparing ASD and control groups across five temporal scales, using standardised sample entropy maps and threshold-free cluster enhancement (TFCE) correction with permutation testing (*p* < .05).

**Figure 3.**
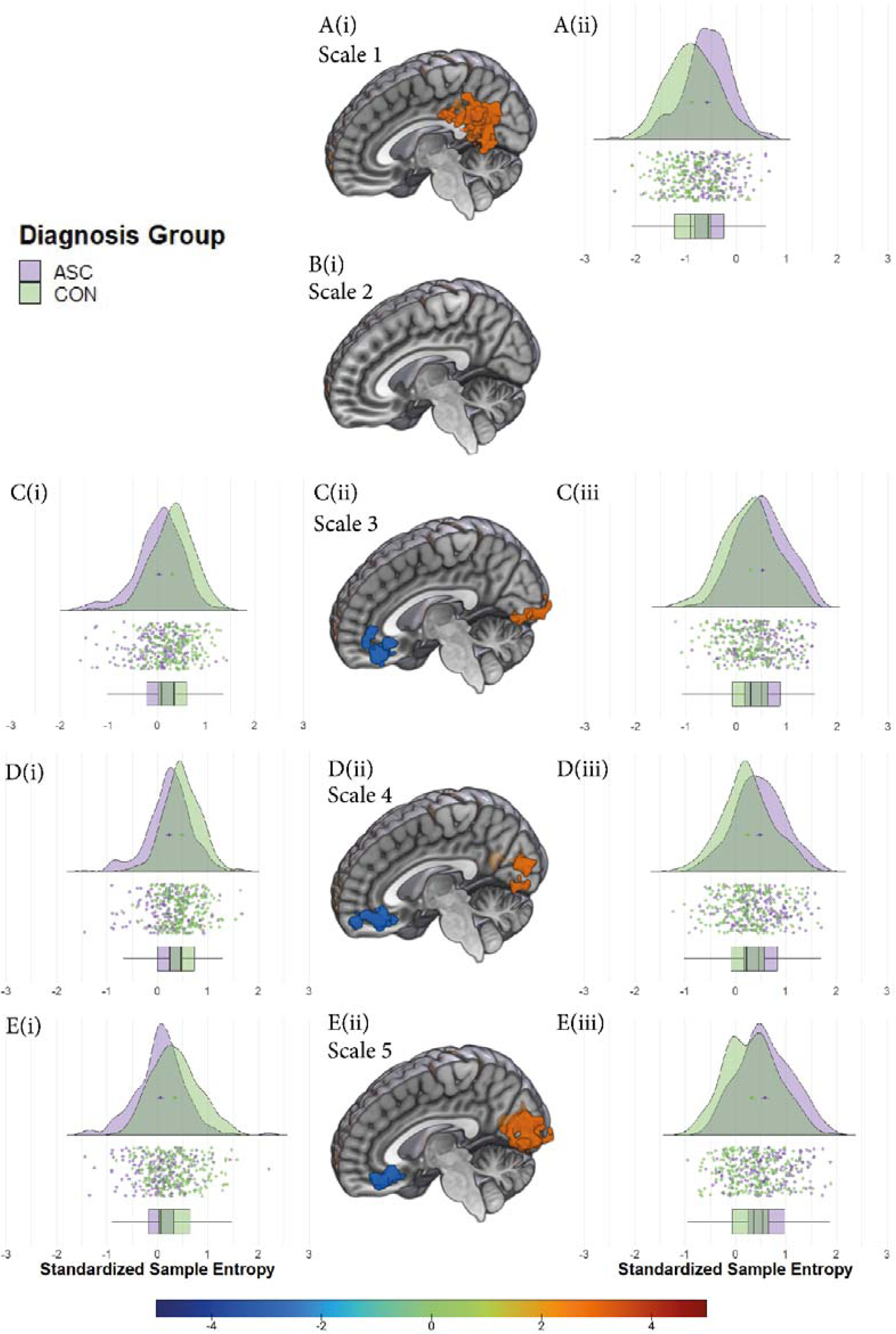
Between-group differences in voxel-wise multiscale sample entropy (MSE) across five temporal scales. For each scale (A–E), sagittal brain slices (middle panels, A–E(ii)) show significant clusters of group differences in entropy between individuals with autism spectrum disorder (ASD) and neurotypical controls (CON), based on voxel-wise two-sample *t*-tests. Warm colours indicate higher entropy in the ASD group, while cool colours indicate higher entropy in the CON group. All results are corrected using threshold-free cluster enhancement (TFCE) with voxel-level permutation testing at *p* < .05. The left and right panels for each scale display corresponding violin and box plots of standardised sample entropy values extracted from significant clusters. Purple represents ASD, and green represents CON. Cluster anatomical labels were identified using xjView (http://www.alivelearn.net/xjview/). Full statistical details for each cluster are reported in the main text.

At scale 1, a significant group difference was observed in the right precuneus cortex (MNI: 9, –54, 18), where the ASD group exhibited increased entropy relative to controls (peak *t* = 5.29; 429 voxels, 216 mm³). No significant group differences were found at scale 2.

At scale 3, two clusters showed significant differences: the left medial frontal cortex (MNI: –9, 33, –3) showed reduced entropy in the ASD group (peak *t* = –4.75; 131 voxels), whereas the left intra-calcarine cortex (MNI: –9, –84, 0) exhibited elevated entropy in ASD (peak *t* = 3.80; 66 voxels). These effects were mirrored at scale 4, with significant group differences in the left paracingulate gyrus (MNI: –12, 48, –3; *t* = –4.14; 112 voxels) and left intra-calcarine cortex (MNI: –15, –75, 3; *t* = 3.60; 100 voxels).

At scale 5, the same regions re-emerged, reinforcing the reliability of these findings. The left paracingulate gyrus (MNI: –12, 45, –6) showed lower entropy in ASD (*t* = – 4.31; 112 voxels), while the left intra-calcarine cortex (MNI: –9, –72, 3) showed higher entropy in ASD (*t* = 4.39; 346 voxels). The spatial consistency of these clusters from scales 3 to 5 highlights robust, scale-dependent group differences in regional brain complexity.

### Results from Voxel-wise Mixed Effect Analysis

A significant main effect of diagnosis was observed in three regions. Lower entropy in the ASD group was detected in the left middle frontal gyrus (MNI: –24, 60, 12; *t* = –4.50; 27 voxels). In contrast, higher entropy in ASD was observed in the left cuneus (MNI: –9, –72, 3; *t* = 4.52; 245 voxels) and the right posterior cingulate cortex (MNI: 6, –54, 24; *t* = 3.54; 22 voxels).

In addition,a significant Group × Time Scale interaction was identified in the right precuneus (MNI: 9, –54, 18; *t* = 5.44), suggesting a scale-dependent divergence in entropy trajectories between groups. Specifically, while both groups exhibited increasing entropy from Scale 1 to Scale 2, only the ASD group showed a subsequent decline at higher scales, whereas entropy in the control group remained relatively stable. The corresponding cluster is visualised in Figure 4B, with group-wise entropy trajectories shown in panel B(iii).

**Figure 4.**
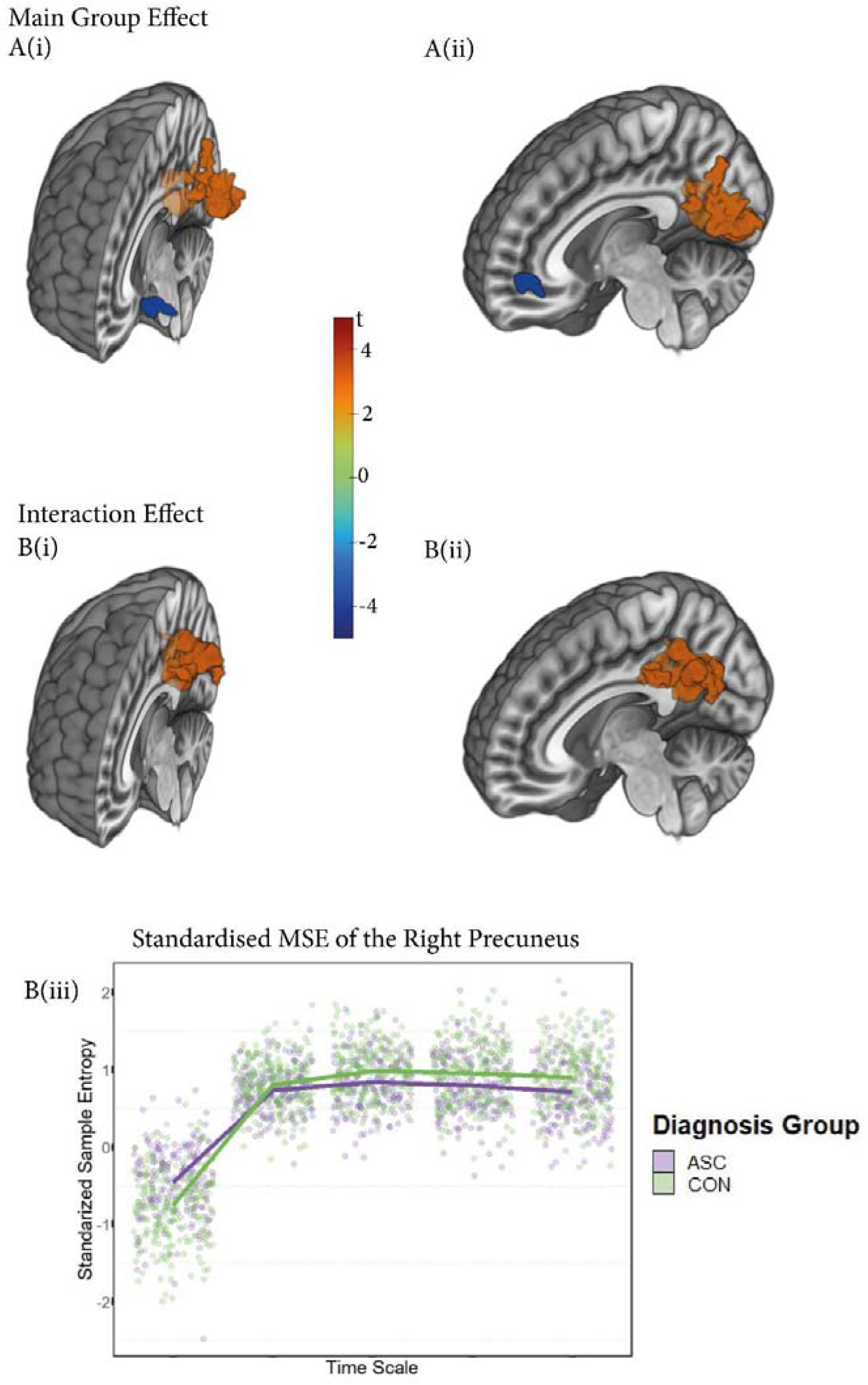
Voxel-wise mixed-effects analysis of multiscale sample entropy (MSE) showing the main effect of the diagnosis group (A(i)–A(ii)) and the Group × Time Scale interaction effect (B(i)–B(ii)). Warm colours indicate higher MSE in the ASD group; cool colours indicate higher MSE in the control group. All results are thresholded using threshold-free cluster enhancement (TFCE) at *p* < .05. Panel B(iii) displays standardised MSE values extracted from the right precuneus cluster (MNI: 9, -54, 18), which exhibited a significant Group × Time Scale interaction (*t* = 5.44). The plot reveals that while both groups show increasing entropy from Scale 1 to Scale 2, the ASD group exhibits a notable decline in entropy across higher time scales, in contrast to the control group whose entropy remains relatively stable. This suggests a scale-dependent reduction in signal complexity in ASD. Purple represents ASD; green represents controls. T-values represent voxel-wise test statistics.

### Exploratory Correlation with ADOS Scores

No significant correlations were found between sample entropy in the significant clusters and ADOS total scores among ASC participants (*p* > .05). Full details of this exploratory analysis, including site distribution and region-specific statistics, are reported in the Supplementary Materials (Supplementary Table 4).

## Discussion

This study investigated differences in resting-state brain signal complexity between individuals with ASD and neurotypical controls, using multiscale sample entropy (MSE) derived from eyes-open fMRI data in a large, multi-site cohort. By applying voxel-wise analysis across multiple temporal scales, we identified robust group-level differences in BOLD signal complexity. Individuals with ASD exhibited increased entropy in posterior regions such as the precuneus and intra-calcarine cortex, and decreased entropy in prefrontal and midline areas including the medial frontal and paracingulate cortices. These effects were especially pronounced at intermediate to coarse time scales, and a significant diagnosis-by-scale interaction in the right precuneus suggested differential temporal profiles of signal irregularity between the two groups. In addition to these voxel-level findings, supplementary analyses at the network level indicated reduced relative entropy in the default mode and frontoparietal networks in ASD, consistent with prior reports of altered large-scale network dynamics in the condition.

Notably, entropy alterations were concentrated within the default mode network (DMN), a set of functionally interconnected regions including the medial prefrontal cortex (mPFC), posterior cingulate cortex (PCC), and precuneus. Prior studies using linear functional connectivity measures have reported inconsistent findings across DMN subregions in ASD, potentially reflecting differential sensitivity to underlying signal dynamics (Lynch et al., 2013; Bathelt & Geurts, 2021). The present findings offer a more nuanced perspective by revealing region-specific changes in signal complexity: the anterior DMN exhibited reduced entropy at coarse scales, while the posterior DMN showed elevated entropy at fine scales followed by a rapid decline. These opposing patterns underscore the heterogeneity of DMN subcomponents and suggest distinct disruptions in temporal organisation within anterior and posterior nodes in ASD (Maximo et al., 2021).

The present analysis identified that individuals with ASD exhibited reduced complexity in the prefrontal cortex, as measured by MSE from resting-state fMRI data. This reduction was most pronounced at coarse time scales, which are thought to reflect slower fluctuations and capture long-range temporal dependencies in neural activity (Vakorin, Lippe, & McIntosh, 2011).This finding suggests a disruption in the integrative temporal organisation of prefrontal dynamics in ASD (Maximo, Nelson, & Kana, 2021). Although sample entropy quantifies signal unpredictability, physiological complexity is not equivalent to randomness; rather, it reflects structured variability across multiple time scales. Crucially, it includes features such as long-range fractal correlations and nonlinear coupling between interacting components (Goldberger, 1996). The observed reduction in prefrontal entropy at coarser scales aligns with the "loss of complexity" hypothesis, which posits that pathological states are characterised by a degradation of hierarchical system organisation and reduced adaptive capacity (Goldberger, 1997).

This interpretation is further supported by studies examining fractal structure in neural and physiological signals, which have consistently shown that a breakdown of long-range correlation properties is associated with impaired regulatory function across health and disease (Courtiol et al., 2016; van Noordt & Willoughby, 2021). According to this view, decreased system complexity can result from either the impairment or loss of functional components, or from disrupted nonlinear coupling between them (Lipsitz, 1992). In the context of ASD, the reduced prefrontal entropy observed at slower temporal scales may therefore reflect disrupted integrative capacity and diminished neural adaptability.

In contrast, participants with ASC showed elevated entropy at the finest time scale in the posterior cingulate cortex (PCC) and precuneus (PCUN), followed by a gradual decline across coarser scales. This pattern indicates that BOLD signals in these regions are highly irregular over short timescales but lack consistent structure over longer temporal windows. Such a pattern suggests a shift toward uncorrelated randomness, characterised by heightened entropy at shorter time scales followed by a decayed entropy at longer time scales (Yang et al., 2015). Such a profile may indicate disorganised or inefficient temporal structure in posterior midline regions relative to anterior areas. Notably, similar scale-dependent shifts in entropy have been observed in an ASD case study following electroconvulsive therapy, where decreased frontal entropy and increased occipital entropy were associated with clinical improvement (Okazaki et al., 2015). These findings raise the possibility that spatially and temporally specific entropy profiles could serve as neurophysiological markers of atypical functional organisation in ASD.

Furthermore, these findings emphasise the importance of investigating multiscale sample entropy (MSE) patterns rather than solely focusing on a single scale. In traditional single-scale entropy studies, there have been reports of both increases and decreases in the complexity of brain signals (Maximo et al., 2021; Zhang et al., 2020). This contradiction is likely caused by sample entropy being initially designed to evaluate the predictability or regularity of a series of data points (Delgado-Bonal & Marshak, 2019) rather than serving as a direct indicator of physiological complexity. Single-scale entropy does not explicitly explore the nonlinear properties of the signal or quantify fractal scaling behaviour, as an increase in sample entropy does not necessarily indicate a corresponding increase in physiological complexity (Vaillancourt & Newell, 2002).

This study provides new insights into how brain signal complexity differs within the DMN between individuals with ASD. The DMN is typically deactivated during externally focused tasks (Fox et al., 2005; Buckner et al., 2008) and activated during rest, particularly when individuals engage in self-referential or introspective mental activity. Accordingly, it has been widely characterised as a core resting-state network (Greicius et al., 2003; Korgaonkar et al., 2014). The observed alterations in DMN complexity in ASC support the view that resting-state activity is not simply passive or task-free, but rather reflects ongoing, internally driven cognitive processes (Biswal, 2012; Buckner et al., 2013). The demonstrated sensitivity of entropy-based features to task-related neural dynamics, even in resting-state data (Lin et al., 2022; Nezafati et al., 2020), further emphasises the value of complexity measures for probing intrinsic brain function.

Voxel-wise one-sample t-test results also revealed systematic differences between cortical and white matter regions across temporal scales. At the finest scale (scale 1), both diagnostic groups showed significantly lower sample entropy in cortical structures compared to the brain-wide average, while white matter regions exhibited relatively higher entropy. As time scale increased, entropy values in cortical regions rose in a pattern consistent with the expected trajectory of complex signals, suggesting the emergence of new temporal patterns in cortical BOLD activity. In contrast, white matter entropy decreased rapidly across scales, displaying an opposite trend to that of grey matter. At scale 1, this elevated white matter entropy may be attributable to increased BOLD signal variability, potentially due to greater signal dispersion from large white matter volumes (Wang et al., 2021). Unlike cortical regions involved in active computation, white matter is primarily composed of myelinated axons that enable long-range communication (Blumenfeld, 2010). The BOLD signal in white matter is often considered to be dominated by noise (Murphy et al., 2013), and the rapid entropy decline across scales likely reflects a lack of structured temporal information. These results provide additional validation for the standardised multiscale entropy approach, demonstrating its capacity to differentiate between grey and white matter based on intrinsic signal properties.

Although we observed clear alterations in DMN complexity in individuals with ASD, these features were not significantly correlated with ADOS scores in the current sample. This may suggest that entropy-based measures capture broader neurophysiological characteristics that are not directly aligned with behavioural symptom severity as measured by diagnostic scales. Rather than diminishing their relevance, this finding points to the potential of complexity features as markers of underlying brain function that may extend beyond symptom-specific expression. Future work will be important to assess the clinical utility and diagnostic specificity of these features. Approaches such as transdiagnostic comparisons and age-normed modelling may offer clearer insights into whether these complexity alterations reflect condition-specific neural signatures or broader patterns of neurodevelopmental divergence (Li et al., 2020; Parkes et al., 2020; Bathelt et al., 2020).

## Limitations

This study has several limitations. First, although we identified group-level differences in brain signal complexity, we did not observe significant correlations with behavioural measures, which limits conclusions about symptom-specific relevance. Second, the case–control design does not account for individual variability within the ASC population or developmental changes over time. Future research may benefit from transdiagnostic comparisons to assess the specificity of these findings, and from age-normed modelling to examine how individuals with ASC deviate from typical developmental trajectories. Finally, although data were drawn from a large multi-site dataset with harmonised preprocessing, residual site-related variability cannot be entirely ruled out.

## Conclusion

This study revealed altered brain signal complexity in individuals with autism spectrum condition, particularly within the default mode network. Using multiscale entropy features from resting-state fMRI, we observed reduced complexity in prefrontal regions and a distinct pattern in posterior midline areas, with increased entropy at fine time scales followed by a decline at coarser scales. These findings underscore the importance of examining brain dynamics across multiple temporal scales and highlight how multiscale entropy captures neural differences that extend beyond what traditional single-scale entropy or connectivity measures can reveal. This approach offers a promising direction for advancing our understanding of intrinsic brain organisation in neurodevelopmental conditions.

