## Supplementary Materials for "Multiscale Entropy of Resting-State fMRI Signals Reveals Differences in Brain Complexity in Autism"

**Supplementary Figure 1**

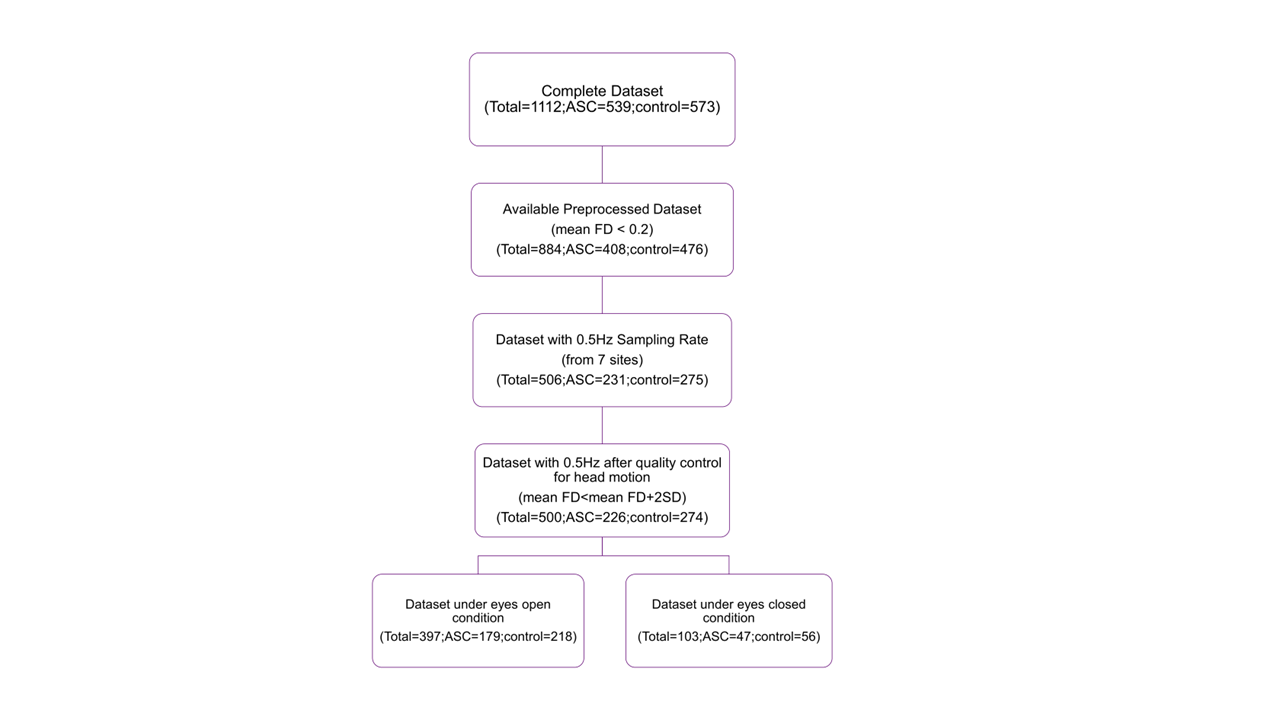

This figure illustrates the data selection procedure used in the current study. The full ABIDE I dataset includes 1,112 participants, of which 539 were diagnosed with autism spectrum disorder (ASD) and 573 were neurotypical controls. The initial selection was based on head motion, where participants with a mean framewise displacement (FD) exceeding 0.2 mm were excluded. From this subset, only those with a sampling rate of 0.5 Hz (i.e., TR = 2000 ms) were retained to ensure uniform temporal resolution for entropy estimation. Participants with excessive motion, defined as mean FD greater than the group mean plus two standard deviations within each site, were further excluded. Finally, only participants scanned under the eyes-open condition were retained for analysis. The resulting dataset consisted of 500 participants, including 226 individuals with ASD and 274 neurotypical controls.

### **Supplementary Figure 2**

*Distribution of Age among ASC and Control groups at the final dataset under eyes-open condition.*
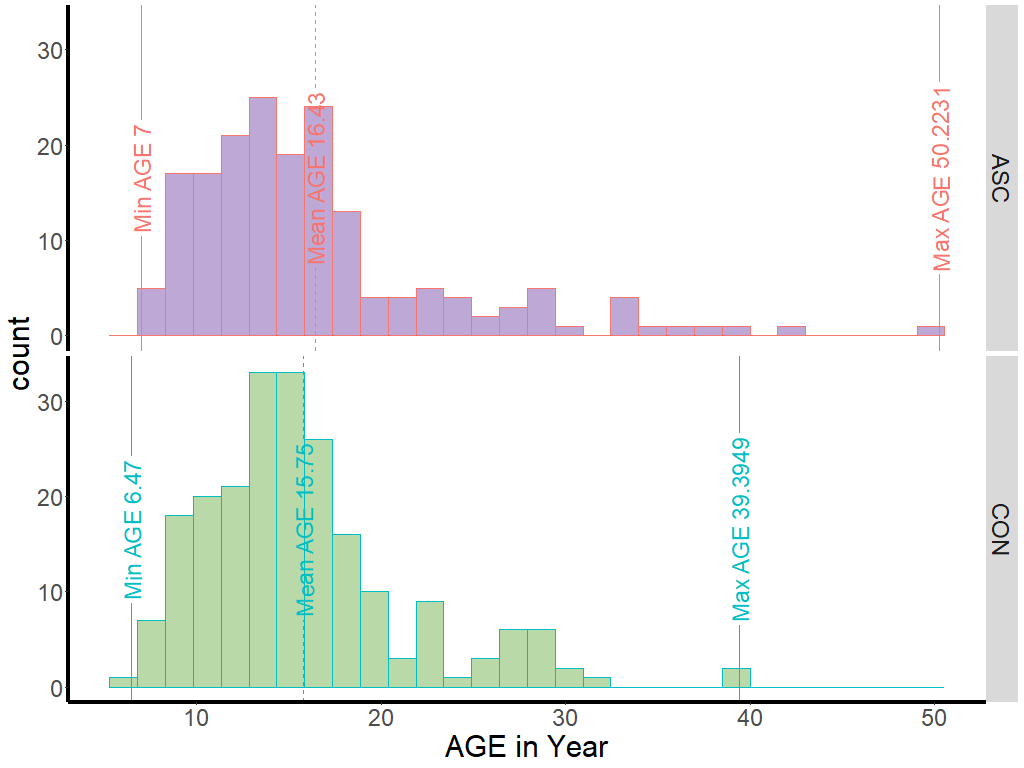

*Note.* Vertical lines indicate minimum, averaged, and maximum age within each group from left to right at each panel.

**Age Distribution in Final Sample**

This figure shows the distribution of ages for participants in the final sample, separated by the diagnostic group. Among participants with ASD, ages ranged from 7.00 to 50.22 years (Mean 16.43 years old, SD 7.17 years old), while controls ranged from 6.47 to 39.39 years (Mean 15.75 years old, SD 5.67 years old). Vertical lines in each histogram represent the minimum, mean, and maximum age for each group. No statistically significant difference in age was observed between groups, *t*(498) = 1.04, *p* = .299.

### **Supplementary Figure 3**

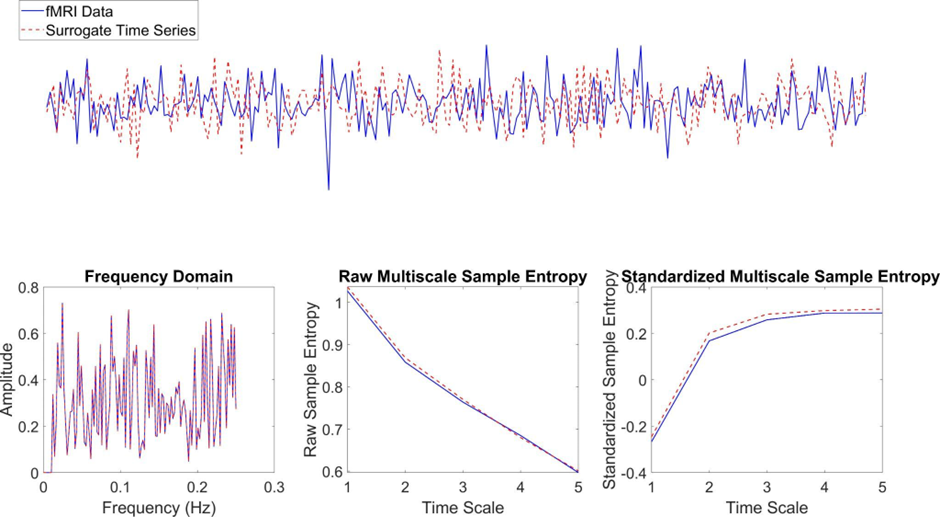

### **Supplementary Figure 4**

*Averaged relative sample entropy.*

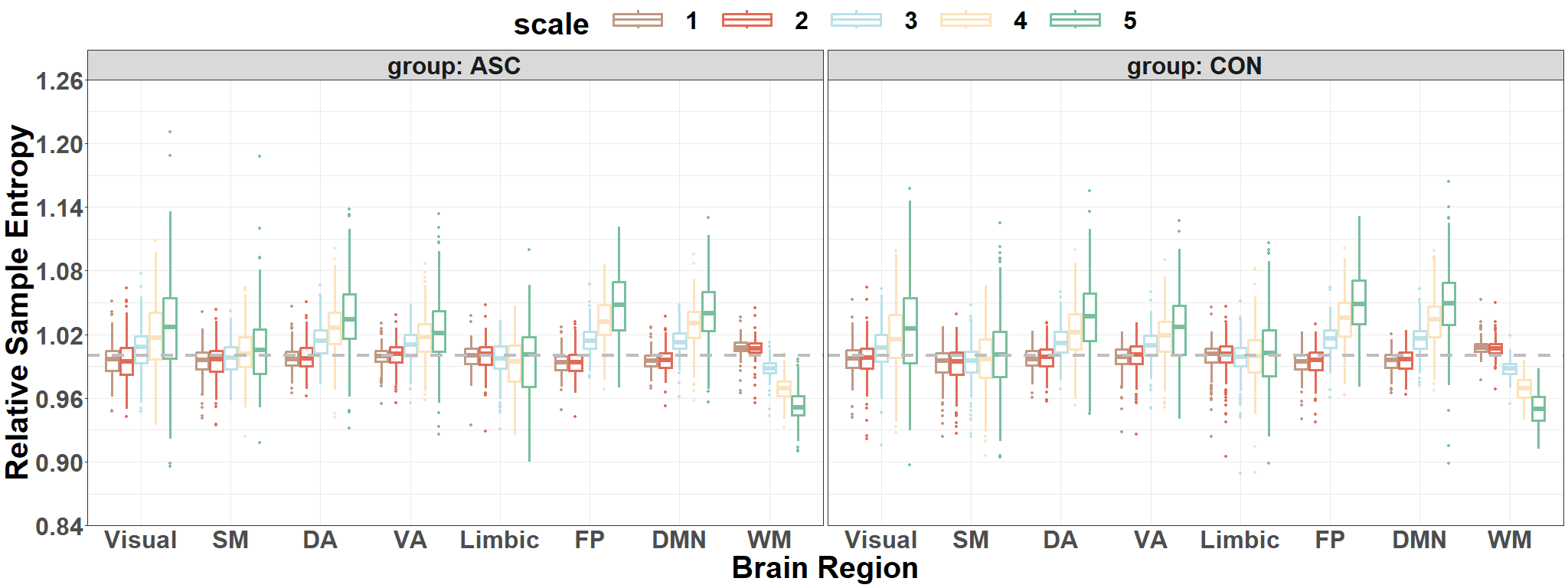

*Note*. Visual = Visual Network; SM = Somatomotor Network; DA = Dorsal Attention Network; VA = Ventral Attention Network; Limbic = Limbic Network; FP = Frontoparietal Network; DMN = Default Mode Network; WM = White Matter.

### **Supplementary Table 1**

**Table 2**

*A comprehensive overview of the acquisition parameters used across different sites in the study*

| Site | Voxel size | Slice thickness | TR | TE | Time points |
| --- | --- | --- | --- | --- | --- |
| NYU | 3.0*3.0*4.0 mm | 4.0 mm | 2000 ms | 15ms | 180 |
| SDSU | - | 3.4 mm | 2000 ms | 30 ms | 180 |
| STANFORD | 3.125*3.125 *4.500 mm |  | 2000 ms | 30 ms | 180 |
| TRINITY | 3.0*3.0*3.0 mm | 3.5 mm | 2000 ms | 28 ms | 150 |
| UM | 3.438*3.438*3.0mm | 3.0 mm | 2000 ms | 30 ms | 300 |
| USM | 3.4*3.4*4.0mm | 3.0 mm | 2000ms | 28ms | 240 |
| YALE | 3.4*3.4*4.0mm | 4.0 mm | 2000ms | 25ms | 200 |

*Note*. TR = repetition time. TE = echo time.

NYU = NYU Langone Medical Center. SDSU = San Diego State University.

STANFORD = Stanford University. TRINITY = Trinity Centre for Health Sciences. UM = University of Michigan. USM = University of Utah School of Medicine. YALE = Yale Child Study Center.

**Acquisition Parameters Across Sites**

Resting-state fMRI data included in the analysis were acquired from seven ABIDE I sites using echo-planar imaging sequences with a repetition time of 2000 ms. Although all sites adhered to this TR, some variability existed in acquisition parameters such as voxel size, slice thickness, echo time (TE), and scan length. In-plane voxel dimensions ranged from 3.0 to 3.4 mm, with slice thicknesses between 3.0 mm and 4.5 mm. Echo times varied from 15 ms to 30 ms, and the number of volumes per scan ranged from 150 to 300 depending on the site. These site-specific acquisition parameters are detailed in Supplementary Table 1 and provide context for the technical consistency of the included data, ensuring that multiscale entropy estimation was applied across comparably sampled datasets.

### **Supplementary Table 2**

*Site-Wise Pre-processed Data Exclusion Criteria Using Head Motion Parameter (FD_Jenkinson)*

| Site | Original  Sample size | Head Motion Criteria  (M += 2SD) | Independent Sample t-tests between Group in Head Motion Parameter | Final Sample Size |
| --- | --- | --- | --- | --- |
| NYU | 184 | 0.085+=0.033 0.068+=0.025 | *t*=3.471, *p*<0.01 | 166 (69 in autism;97 in control) |
| SDSU | 36 | 0.070+=0.051  0.072+=0.036 | *t*=-0.084, *p*=0.934 | 33 (12 in autism;21 in control) |
| STANFORD | 40 | 0.117+=0.032  0.124+=0.062 | *t*=-0.395, *p*=0.696 | 36 (17 in autism;19 in control) |
| TRINITY | 49 | 0.114+=0.046  0.082+=0.019 | *t*=2.918, *p*<0.01 | 44 (21 in autism;23 in control) |
| UM_1 | 110 | 0.104+=0.048  0.075+=0.033 | *t*=3.123, *p*<0.01 | 82 (36 in autism;46 in control) |
| UM_2 | 35 | 0.101+=0.066  0.092+=0.093 | *t*=0.300, *p*=0.766 | 31(12 in autism;19 in control) |
| USM | 101 | 0.107+=0.050  0.107+=0.046 | *t*=-0.006, *p*=0.995 | 60 (37 in autism;23 in control) |
| YALE | 56 | 0.104+=0.050  0.088+=0.043 | *t*=1.189, *p*=0.241 | 48 (22 in autism;26 in control) |

*Note*. NYU = NYU Langone Medical Center. SDSU = San Diego State University.

STANFORD = Stanford University. TRINITY = Trinity Centre for Health Sciences. UM = University of Michigan. USM = University of Utah School of Medicine. YALE = Yale Child Study Center.

**Head Motion Quality Control by Site**

To control for motion-related artefacts, mean framewise displacement was calculated using Jenkinson’s method. Participants whose mean FD exceeded the group mean plus two standard deviations were excluded on a per-site basis. Final sample sizes, site-level means, thresholds, and independent-sample t-test results comparing FD between diagnostic groups are reported in Supplementary Table 2. In most sites, there were no statistically significant differences in mean FD between the ASD and control groups. Exceptions included NYU, Trinity, and UM_1, where group differences were statistically significant, though the effect sizes were modest and the retained samples were well within conservative motion thresholds. These procedures ensured that motion-related confounds were minimised in the final analysis.

### **Supplementary Table 3**

### **Available Samples with ADOS Scores across Different Sites**

| **Site** | **Sample Size** | **ADOS Mean (SD)** |
| --- | --- | --- |
| NYU | 60 | 11.433 (4.323) |
| SDSU | 11 | 10.545 (4.321) |
| USM | 37 | 12.838 (3.023) |

Note.
 NYU = NYU Langone Medical Center.
 SDSU = San Diego State University.
 USM = University of Utah School of Medicine.

**Supplementary Table 4**

**Correlation Between Entropy in Significant Clusters and ADOS Scores**

| **Brain Region** | **Time Scale** | **r** | **p** |
| --- | --- | --- | --- |
| Right precuneus cortex | 1 | 0.061 | 0.533 |
| Left medial frontal cortex | 3 | 0.062 | 0.523 |
| Left intra-calcarine cortex | 3 | -0.118 | 0.226 |
| Left paracingulate gyrus | 4 | 0.062 | 0.523 |
| Left intra-calcarine cortex | 4 | -0.127 | 0.190 |
| Left paracingulate gyrus | 5 | 0.002 | 0.987 |
| Left intra-calcarine cortex | 5 | -0.101 | 0.296 |

Note. ADOS = Autism Diagnostic Observation Schedule total score.
 Pearson’s correlation was used to assess the relationship between relative sample entropy values from significant clusters and ADOS scores among ASC participants. No statistically significant correlations were observed.

### **Supplementary Methods**

**Rationale for Restricting the Dataset to TR = 2000 ms**

Multiscale entropy (MSE) relies on a coarse-graining process that involves averaging adjacent data points in the time series to generate signals at progressively longer temporal scales. The temporal interpretation of each scale is directly tied to the repetition time (TR) of the data. Inconsistencies in TR across participants would result in the same coarse-graining scale representing different real-world durations, thus confounding any group-level or across-subject comparisons. For example, a scale factor of 5 corresponds to 10 seconds at TR = 2000 ms but 15 seconds at TR = 3000 ms. Additionally, longer TRs yield shorter time series, which limits the number of usable scales and reduces the reliability of entropy estimates. While interpolating or resampling time series might superficially harmonise sampling rates, these methods introduce temporal smoothing and artefactual dependencies, which are detrimental to the estimation of neural signal complexity.

Previous studies have recommended using time series of at least 100–200 points for stable entropy estimation (Sokunbi, 2014; Courtiol et al., 2016). To ensure the physiological validity of each coarse-graining scale, and to preserve consistency across the sample, we included only participants whose data were acquired with a TR of 2000 ms, corresponding to the most common and well-sampled subset within the ABIDE I dataset. This allowed for the estimation of multiscale entropy without the need for resampling or interpolation, thereby ensuring methodological rigor and comparability across participants.

#### **Entropy Computation and Implementation Details**

**Mathematical Definition of MSE**

***Multi-Scale Entropy***

Multi-Scale sample entropy (MSE) computes the sample entropy on the original signal and on coarse-grained time series that are derived from the original signal. MSE calculation incorporates two procedures in each epoch and for each electrode independently. Firstly, the algorithm generates coarse-grained time series by progressively downsampling the time series$\left\{ \chi₁,\chi₂,\ldots\chiɴ \right\}$. As for time scale factor 𝜏, each element $\left( i \right)$ from the coarse-grained time series $\left\{ Y\tau\left( i \right) \right\}$is obtained by averaging 𝜏 consecutive data points, resulting in a time series with a length of exactly divisible by 𝜏 into the original length. Specifically, the time series associated with scale factor 1 is the original data and scale factor 2 is the average of consecutive pairs of data points and so forth for increasing scales. As such, the element of a coarse-grained time series $i$ is calculated according to:

$\left\{ Y\tau\left( i \right) \right\}=\frac{1}{\tau}\sum_{i=1}^{\frac{N}{\tau}} \left( \chi{}_{((i-1)*\tau+1)}: \chi{}_{(i*\tau)} \right)$	(Equation 1)

where $N$ is the length of the original signal. Second, the algorithm computes the sample entropy for each coarse-grained time series. Sample entropy is defined by the negative natural logarithm of the conditional probability that a time series of length $\frac{N}{\tau}$, having repeated itself within a tolerance $\gamma$(similarity threshold) for $m$ points (the length of sequences to be compared), will also repeat itself for $m+1$ points, without allowing self-matches. The pattern length $m$ was fixed to 2 and the similarity factor $\gamma$ was set to 0.60 * standard deviation of the time series in this study; that is, data points were considered to be indistinguishable if the absolute amplitude difference between them was ≤ 60% of the standard deviation of the time series.

Before the SE calculation, all time series were centred and normalised to standard deviation 1 to avoid the bias of varying range of amplitude across channels and datasets. SE is proposed to calculate the distance between any pairwise elements and count the number of times when this distance is under the threshold $\gamma$. To do this, for a time series with $N$ elements, forms a $(N- m+1)$ vectors $U_{m} (i)$ $\left| 1<i<N- m+1 \right|$,where $U_{m} (i)=\chi(i+(m-1))$ is the vector $m$ points from $\chi_{(i)}$to $\chi_{(i +(m-1))}$. Then the built-in matlab function *pdist* is used to determine the pairwise Chebyshev distance $d\left[ U_{m} (i) -U_{m} (j) \right]\left( i\neq j \right)=max\left\{ {|U}_{m} (i)-U_{m} (j)| \right\}$, then it is extended to the$m+1$. The number of thresholded pairwise distances are counted respectively. The quantity is:

$A_{i}^{m}(\gamma)=\sum_{i=1\left( i\neq j \right)}^{N-m} (d\left[ U_{m} (i) -U_{m} (j) \right]<\gamma)$ and$B_{i}^{m+1}(\gamma)=\sum_{i=1\left( i\neq j \right)}^{N-m+1} (d\left[ U_{m+1} (i) -U_{m+1} (j) \right]<\gamma)$

(Equation 2)

and sample entropy equation is:

$Sample Entropy(m, \gamma, N) = -\frac{N-m+1}{N-m-1}*log(\frac{A(m, \tau)}{B(m+1, \tau)})$	(Equation 3)

**Validation Using Surrogate Data**

To assess the sensitivity of MSE to nonlinear temporal structure in the fMRI signal, we conducted a control analysis comparing entropy profiles from real BOLD time series to phase-randomised surrogate data. Although the two signals shared similar power spectra, MSE diverged at coarser scales, indicating that the observed entropy captures nonlinear dependencies not preserved in the surrogate data. Results are visualised in Supplementary Figure 3.

**Validation of Multiscale Sample Entropy on fMRI Data Using Phase-Shuffled Surrogates**

To further examine the sensitivity of multiscale sample entropy (MSE) to nonlinear temporal dependencies in resting-state fMRI signal irregularities and verify its validity, phase-shuffled surrogate time series were introduced. These surrogate time series were generated by subjecting the original fMRI signals to a Fourier transform (FFT), uniformly randomly shuffling the phase of the Fourier components, and then applying the inverse Fourier transform to reconstruct the time domain. To enhance reliability in identifying nonlinearity, the iterated amplitude-adjusted Fourier transform (IAAFT) technique was employed to shift the nonlinear correlation structure (multifractality) while keeping the degree of linear correlation (persistence) and the amplitude distribution (Theiler et al., 1992; Schreiber & Schmitz, 1996). This procedure includes the following steps:

1. Perform FFT on the original time series.
2. Initialise the surrogate time series and perform FFT on the surrogate time series.
3. Match the amplitude of the original FFT and surrogate FFT.
4. Generate random phase shifts.
5. Apply the phase shifts to the amplitude-matched FFT.
6. Apply inverse FFT to obtain the phase-shuffled surrogate time series.

The phase-shuffled surrogate time series were obtained through a maximum of 100 iterations. The entire execution was carried out using a custom MATLAB script available on GitHub at the following link: <https://github.com/wenyixiao0058/MSE_effectiveness.git>, specifically referring to the *"phase_suffled_surrogate.m"* file, which employs built-in functions like fft for extracting FFT characteristics and ifft for reversing the data to its original time domain representation.

Supplementary figure 3 shows an example using a resting-state fMRI signal. The upper panel depicts a time series from a random grey matter voxel (solid blue line) alongside a phase-shuffled surrogate time series (dashed red line). Notably, the power spectrum of both the original fMRI and phase-shuffled time series is identical. As the power spectrum reflects a linear process but lacks phase information, the phase-shuffled time series is expected to exhibit higher entropy than the original fMRI time series due to its increased irregularity.

MSE assigns greater entropy to the phase-shuffled surrogate time series than the original time series, indicating sensitivity to nonlinear dynamics. In this case, the decreasing trend in MSE for the fMRI signal may reflect insufficient data points, residual noise artefacts, or limitations in capturing frequency components due to filtering methods employed in this investigation. A detailed exploration of these factors exceeds the scope of the current study but could be pursued in future research.

The current study adopts standardised multiscale sample entropy for further analysis and interpretation. This approach effectively captures enhanced nonlinear features of the phase-shuffled surrogate time series and reproduces key MSE findings, including group and condition differences, in subsequent analyses.

**Spatial Normalisation and Relative Entropy (smentropy)**

To facilitate comparison of local entropy relative to each participant’s overall brain-wide signal complexity, voxel-wise multiscale entropy (MSE) values were normalised by the global mean entropy for each subject. This yielded individual *smentropy* maps representing relative entropy, where a value of 1 indicates a voxel with entropy equal to the participant’s mean, and values above or below 1 reflect proportionally higher or lower regional entropy, respectively. Global mean entropy was computed within a common group-level mask containing voxels present in at least 90% of participants.

Voxel-wise one-sample *t*-tests were performed across participants within each diagnostic group (ASD and controls separately) to evaluate whether *smentropy* significantly deviated from 1 at each voxel. These tests were implemented in DPABI and corrected for multiple comparisons using threshold-free cluster enhancement (TFCE). TFCE-derived *t*-maps were transformed to *Z*-maps using the y_TFRtoZ function from the DPABI Statistical Analysis module (<https://github.com/Chaogan-Yan/DPABI/tree/master/StatisticalAnalysis>). Results were visualised with MRIcroGL (Rorden & Brett, 2000).

**First-Level Analysis: Time Series Extraction and Entropy Estimation**

Voxel-wise multiscale sample entropy (MSE) was calculated from resting-state fMRI data for each participant in the ASD and control groups. Data were selected from seven ABIDE I sites (NYU, SDSU, Stanford, Trinity, UM1&2, USM, and Yale), each with a consistent temporal resolution of 0.5 Hz (TR = 2000 ms). Due to observed variability in signal amplitude across sites, all voxel-wise BOLD time series were normalised to unit standard deviation prior to entropy estimation to ensure comparability.

Multiscale sample entropy (MSE) maps were computed in template (MNI) space for each participant from the seven included ABIDE I sites. Preprocessed resting-state fMRI data and individual anatomical brain masks were read using MATLAB functions from the SPM toolbox (spm_vol, spm_read_vols). Voxel-wise BOLD time series were extracted within each participant's mask, and MSE was calculated using the custom script multiscale_entropy4fmri.m, generating five entropy values per voxel corresponding to time scales τ = 1 to 5.

To prepare maps for group-level analysis, resulting entropy volumes were smoothed with a 6 mm full-width at half-maximum (FWHM) Gaussian kernel. Brain masks were also warped and smoothed to align with the MNI space. Final entropy maps were saved in NIfTI format using spm_write_vol.

The complete data analysis process, encompassing tasks such as data input, time series extraction, multiscale sample entropy computation, generation of resultant entropy brain maps, and storage of outcomes for individual participants, was executed using the High-Performance Computing resources provided by the University of Sheffield's ShARC (Sheffield Advanced Research Computer) infrastructure. Scripts used in the pipeline are available at: <https://github.com/wenyixiao0058/fmristudy>, including getfmriDat.m for data input, setfmriAggregate.m for output organisation, and the entropy computation script.

**Second-Level Analysis: Group-Level Comparisons**

At the group level, voxel-wise two-sample t-tests were performed to compare MSE values between participants with ASD and neurotypical controls. These tests produced statistical maps indicating voxel-wise group differences across each of the five time scales. The null hypothesis in each case was that there is no difference in entropy values between groups at a given voxel.
When inferring the fMRI spatial content, hundreds of thousands of t-tests were conducted for each voxel simultaneously, which inflates the false positives. Bonferroni correction is too conservative as it considers these t-tests independent, which is different for fMRI inference as these t-tests are related to their neighbouring test. To better correct the multiple comparison problems raised by a mass of t-tests conducted in this voxel-wise study, a permutation test with Threshold-Free Cluster Enhancement (TFCE) was applied to achieve a balance between test-retest reliability and family-wise error rate (under 5%) as well as cluster-based inference which assesses the surprising spatial extent (i.e., the size of cluster connected by voxels). This step was achieved by the DPARSF toolbox integrating PALM(Permutation Analysis of Linear Models) (Winkler et al., 2016). The number of permutations was set to 5,000 using the “no acceleration” method, which is recommended for spatial statistics and provides strong control over the family-wise error rate (FWER) at 5%.

The analysis also highlighted that using non-uniform individual brain masks when computing voxel-wise entropy can introduce false positives in group comparisons. This issue arises when a voxel is present in the brain mask of one participant but absent in another. During the voxel-wise two-sample t-test, these inconsistencies result in comparisons between valid entropy values and missing data, particularly around the cortical boundaries, leading to spurious statistical differences. To avoid this artefact, it is recommended to apply a consistent group-level brain mask across all participants, groups, and sites.

**One sample voxel-wise t-test within each diagnosis group**

To reduce the influence of global inter-participant variability in entropy, as done in other temporal analyses in rs-fMRI study (Li et al. 2021), voxel-wise entropy values were normalised by each individual's global mean entropy, resulting in a relative entropy measure referred to as *smentropy*. This global mean was computed for each participant using a group-level mask derived from the average of all individual brain masks, thresholded at 90% coverage to ensure consistent spatial sampling. The relative entropy value at each voxel represents the ratio of its raw entropy to the global brain average, thereby reflecting regional deviations from overall brain complexity.

Subsequent voxel-wise one-sample t-tests were applied to determine whether regional relative entropy significantly deviated from 1, separately for the ASD and control groups.

These tests were conducted at each of the five temporal scales using the y_TTest1_Image function from the DPABI toolbox (Yan et al., 2016) at each time scale (1-5) respectively.

The resulting t-maps were corrected for multiple comparisons using threshold-free cluster enhancement (TFCE), and then converted into Z-statistic maps using the y_TFRtoZ function. Visualisation of these maps was performed using MRIcroGL (Rorden & Brett, 2000), centred at the MNI coordinates (0.196167, 2.06932, 26.3201 mm). In the resulting figures, yellow-to-red areas indicate regions with significantly higher entropy relative to the global mean, while green-to-blue areas denote regions with significantly lower relative entropy.

Additionally, two-sample t-tests were used to assess group differences in global mean entropy across each time scale. No significant differences in global entropy were found between ASD and control groups (Scale 1: t = -1.111, p = .267; Scale 2: t = -2.532, p = .0117; Scale 3: t = -1.691, p = .092; Scale 4: t = -0.925, p = .355; Scale 5: t = -0.748, p = .455). These findings support the use of normalised *smentropy* maps for comparing spatial entropy distributions across diagnostic groups.

**Two-Sample Voxel-Wise T-Test Across Diagnostic Groups**

To perform group comparisons, individual voxel-wise entropy maps were standardised by subtracting each participant’s global mean and dividing by the standard deviation, resulting in z-scored entropy maps. This standardisation serves two primary purposes: (1) it reduces the impact of inter-subject variability in overall entropy magnitude, and (2) it restores the multiscale sample entropy profile of the fMRI signal, and its validity was confirmed in the pilot test. Furthermore, it effectively discriminates between grey and white matter by capturing their distinct nonlinear characteristics through standardised multiscale sample entropy.

Group-level analyses were then conducted using both voxel-wise two-sample t-tests (y_TTest2_Image) and voxel-wise mixed-effects models (y_MixedEffectsAnalysis_Image) implemented within the DPABI toolbox (Yan et al., 2016) and are publicly accessible at (https://github.com/Chaogan-Yan/DPABI).

The two-sample t-tests evaluated group differences at each entropy scale, while the mixed-effects models examined the scale-wise change in entropy across groups. All voxel-wise statistical tests were corrected for multiple comparisons using a permutation-based TFCE approach, implemented through the PALM (Permutation Analysis of Linear Models) toolbox (Winkler et al., 2016) integrated in DPABI. The number of permutations was set at 5,000, with the “no acceleration” setting selected to ensure robustness for spatial statistics. This correction strategy accounts for spatial dependencies in the data, providing a balance between family-wise error rate control and test–retest reliability.

**Cortical Network-Level Entropy Extraction**

To visualise network-level changes in brain signal complexity, entropy values were extracted from seven cortical networks identified by Yeo et al. (2011), as well as from the white matter mask. The seven canonical networks include: Visual network: Areas in the occipital and parietal lobes involved in processing visual information. Somatomotor network: Frontal and parietal regions involved in movement and sensation. Dorsal attention network: Parietal and frontal areas associated with spatial attention control. Ventral attention network: Frontal and parietal regions involved in non-spatial attention control. Limbic network: Medial temporal lobe, cingulate cortex, and insula, linked to emotional processing and memory. Frontoparietal network: Frontal and parietal areas related to cognitive control and decision-making. Default mode network: Medial prefrontal cortex and posterior cingulate cortex involved in self-referential thinking and mind-wandering.

The cortical network masks were downloaded from Yeo 2011 atlas (<https://surfer.nmr.mgh.harvard.edu/fswiki/CorticalParcellation_Yeo2011>), projected into MNI152 space, and resliced into 3 mm isotropic voxels. Each mask was intersected with the group-level brain mask to ensure coverage overlap. The final masks were applied to each participant’s relative sample entropy maps, and mean entropy was extracted from each network. This process was repeated for both the ASD and neurotypical control groups. The resulting relative sample entropy values were visualised using boxplots (see Supplementary Figure 4).
